# Trade-off between competition ability and invulnerability to predation in marine microbes; protist grazing versus viral lysis effects

**DOI:** 10.1101/2021.10.09.463787

**Authors:** Jinny Wu Yang, Feng-Hsun Chang, Yi-Chun Yeh, An-Yi Tsai, Kuo-Ping Chiang, Fuh-Kwo Shiah, Gwo-Ching Gong, Chih-hao Hsieh

## Abstract

Trade-offs between competition ability and invulnerability to predation are important mechanisms explaining how predation promotes bacterial diversity. However, existence of these trade-offs has apparently not been investigated in natural marine bacterial communities. Here, we address this question with growth-based measurements for each marine bacterial taxon by conducting on-board dilution experiments to manipulate predation pressure and using high-throughput sequencing to assess the response of bacterial communities. We determined that bacterial taxa with a higher predation-free growth rate were accompanied with higher predation-caused mortality, supporting existence of competitiveness-invulnerability trade-off. This trade-off was stronger and more consistent under viral lysis than protist grazing. In addition, predation generally flattened out the rank-abundance distribution and increased the evenness and richness of the bacterial community. These findings supported the “Kill-the-Winner” hypothesis. All experiments supported a significant competitiveness-invulnerability trade-off, but there was substantial variation among bacterial communities in response to predation across experiments conducted in various sites and seasons. Therefore, we inferred that the Kill-the-Winner hypothesis is important but likely not the only deterministic mechanism explaining how predation shapes bacterial assemblages in natural marine systems.

## Introduction

Marine microbes, the foundation of marine food-webs, dominate biogeochemical cycles and regulate nutrient and energy flows in marine systems [1, 2, 3, 4, 5]. Also, microbial diversity is widely suggested as a critical index reflecting or determining marine ecosystem functioning [3, 5, 6]. Mechanisms shaping marine microbial diversity have been intensively studied, with an emphasis on abiotic mechanisms such as environmental productivity or hydrological factors [7, 8, 9]. Biotic mechanisms involving interactions within a community or across trophic levels are critical in shaping marine microbial diversity [10, 11, 12] but are relatively less explored.

Niche trade-off, one of the most common biotic mechanisms essential for species coexistence [13, 14, 15, 16, 17], occurs when species specialize in a certain trait at the cost of another trait. Characterizing trade-offs in microbial species gives insights into mechanistic understandings of their community organization. However, our understanding is mostly confined to the trade-off between species’ ability to acquire various nutrients [17, 18]. Here, we view trade-off in another context, namely the trade-off between species’ competition ability and their invulnerability to predation (hereafter termed competitiveness-invulnerability trade-off).

In microbes, the competitiveness-invulnerability trade-off is more widely known as the Kill-the-Winner hypothesis: competitively superior prey, the “winners,” have higher mortality caused by predation so that inferior competitors can escape from competition exclusion, thereby promoting species coexistence [15]. Competitive superiority can come from species’ high growth rates and/or their ability to maintain high abundance. For example, bacteria with high density of certain cell-surface receptors can also have higher cellular functions promoting rapid growth rate, but increased susceptibility to viral infection [19, 20]. In addition, developing growth-competitive mechanisms would come at the cost of low predation avoidance or defensive mechanisms due to genetic constraints [15, 21]. The competitiveness-invulnerability trade-off, or Kill-the-Winner hypothesis, is a mechanism to explain how predation can promote microbial diversity in the field [21, 22, 23, 24, 25]. However, there is apparently a lack of evidence in natural systems to demonstrate its occurrence and how it may affect marine bacterial diversity.

In this study, we describe the competitiveness of bacteria as their predation-free growth rate (growth-based competitiveness) and initial density (density-based competitiveness). Invulnerability to predation is regarded as a generic term including protist grazing and viral lysis, the major cause of mortality and selection forces affecting bacterial community diversity and composition [25, 26, 27, 28, 29]. Specifically, we coupled dilution experiments and high-throughput sequencing techniques to estimate the predation-free growth rate and the growth rate under protist grazing and viral lysis for each bacterial taxon. With these estimates, we tested the first hypothesis that bacteria’s competitiveness is negatively associated with their invulnerability to predation due to the occurrence of competitiveness-invulnerability trade-off in marine bacterial communities (Hypothesis I). Based on this hypothesis, we expect the bacterial taxon with a higher growth-based competitiveness, i.e., predation-free growth rate, would have higher mortality caused by protist grazing (Prediction 1) or viral lysis (Prediction 2). Furthermore, we also expect the bacterial taxon with higher density-based competitiveness, i.e., initial density, would have higher mortality caused by protist grazing (Prediction 3) and viral lysis (Prediction 4).

Our second hypothesis is that due to the occurrence of competitiveness-invulnerability trade-off, stronger predation pressure increases bacterial diversity (Hypothesis II). This is because predation should alleviate competitive exclusion by suppressing highly competitive species. To test this hypothesis, we examined how bacterial rank-abundance distribution, evenness and richness responded to increasing predation pressure, namely protist grazing, viral lysis, or both. Under the occurrence of competitiveness-invulnerability trade-off (where Hypothesis I is supported), we expected that higher predation pressures lead to a flattened rank-abundance distribution, higher evenness (lower dominancy by a small number of species in the community; Prediction 5) and higher richness (Prediction 6) of the bacterial community.

## Results

### Competitiveness-invulnerability trade-off in marine bacterial community

The bacterial predator-free growth rate was negatively correlated with its invulnerability to predation, regardless of under protist grazing or viral lysis, as shown in LMM analysis (both *p* < 0.001; Table 1). These results supported the occurrence of competitiveness-invulnerability trade-off driven by protist grazing and viral lysis, respectively (Prediction 1 and 2 supported). In addition, this trade-off was stronger under viral lysis than protist grazing (with LMM estimates regression slope = −0.11 and −0.027, respectively). The conclusion was generally the same when examining each experiment separately: Only five of seven experiments had a significant growth-based competitiveness-invulnerability trade-off under protist grazing; in comparison, this trade-off was apparent in all seven experiments when under viral lysis (Figure 1). The results of the permutation test also supported our conclusion (Table S3). Our findings indicated that viral lysis was a stronger and more consistent driving force on growth-based competitiveness-invulnerability trade-off than protist grazing.

**Table 1.** Results of linear mixed-effect model analysis for competitiveness-invulnerability trade-off and predation effects on bacterial community indices, with experiments as the random effect. Bold numbers indicate significant (*p* < 0.05) results.

| Independent variable | Dependent variable | Estimate | Std. Error | *p* value |
| --- | --- | --- | --- | --- |
| **Competitiveness-invulnerability trade-off** |  |  |  |  |
| Competitiveness | Invulnerability |  |  |  |
| Growth-based competitiveness | to protist grazing | -0.027 | 0.002 | **<0.001** |
|  | to viral lysis | -0.114 | 0.003 | **<0.001** |
| Density-based competitiveness | to protist grazing | 0.139 | 0.008 | **<0.001** |
|  | to viral lysis | -0.047 | 0.011 | **<0.001** |
| **Predation effect on bacterial community** |  |  |  |  |
| Predation effect | Community index |  |  |  |
| Protists+viruses | RAD decay coefficient | -0.107 | 0.025 | **<0.001** |
| Protists |  | -0.032 | 0.033 | 0.335 |
| Viruses |  | -0.074 | 0.029 | **0.012** |
| Protists+viruses | Evenness | 0.037 | 0.009 | **<0.001** |
| Protists |  | 0.013 | 0.011 | 0.237 |
| Viruses |  | 0.023 | 0.010 | **0.024** |
| Protists+viruses | Richness | 21.993 | 8.026 | **0.009** |
| Protists |  | 31.051 | 10.159 | **0.004** |
| Viruses |  | -9.831 | 8.761 | 0.264 |

**Figure 1.**
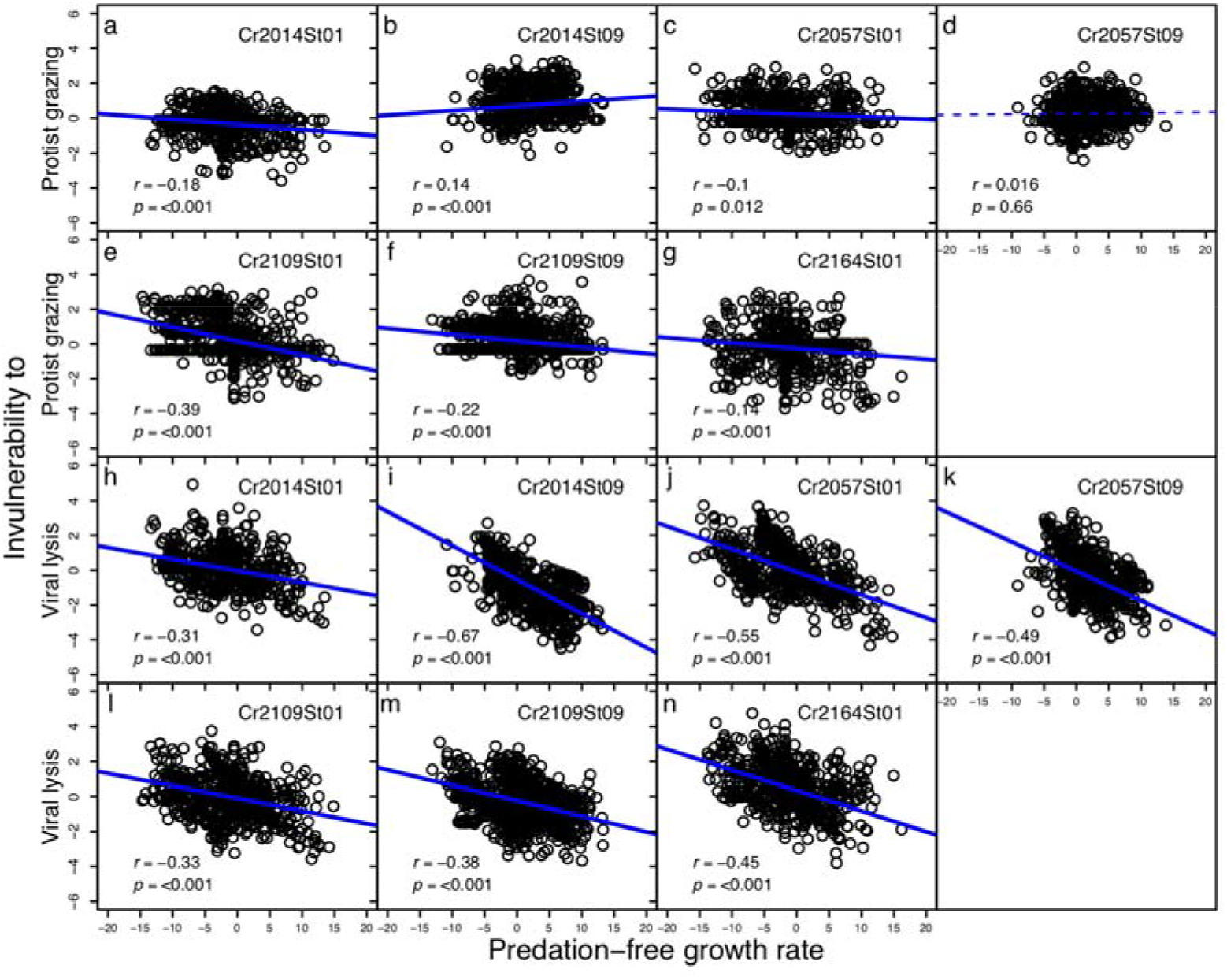
Relationships between predation-free growth rate (hour^−1^) and invulnerability to protists grazing or to viral lysis (hour^−1^), respectively, in each experiment. Values are the Pearson’s correlation coefficient (*r*) and its p value (*p*). Solid and dashed lines indicate significant (*p* < 0.05) and non-significant (*p* > 0.05) linear regression, respectively.

On the contrary, we did not identify a consistent relationship between density-based competitiveness, i.e., initial density, and invulnerability to predation. LMM analysis across seven experiments indicated that bacterial initial density exhibited a significant positive correlation with the invulnerability to protist grazing (*LMM-p* <0.001, Table 1). However, when examining each experiment separately under protist grazing, there were only positive and no relationships (Figure 2 and Figure S4). Therefore, a density-based competitiveness-invulnerability trade-off under protist grazing was not observed in our system (Prediction 3 not supported). Despite a significantly negative correlation between density-based competitiveness and invulnerability to viral lysis in LMM (*LMM-p* <0.001) (Table 1), only three sets of experiments had the consistent and weak relationship (Pearson’s correlation coefficient =−0.07, −0.17, and −0.14, *p* value = 0.04, <0.001 and <0.001, for Figure 2h, 2i and, 2m, respectively). Therefore, we were not able to make a clear conclusion of the existence of this density-based competitiveness-invulnerability trade-off under viral lysis (Prediction 4 not supported).

**Figure 2.**
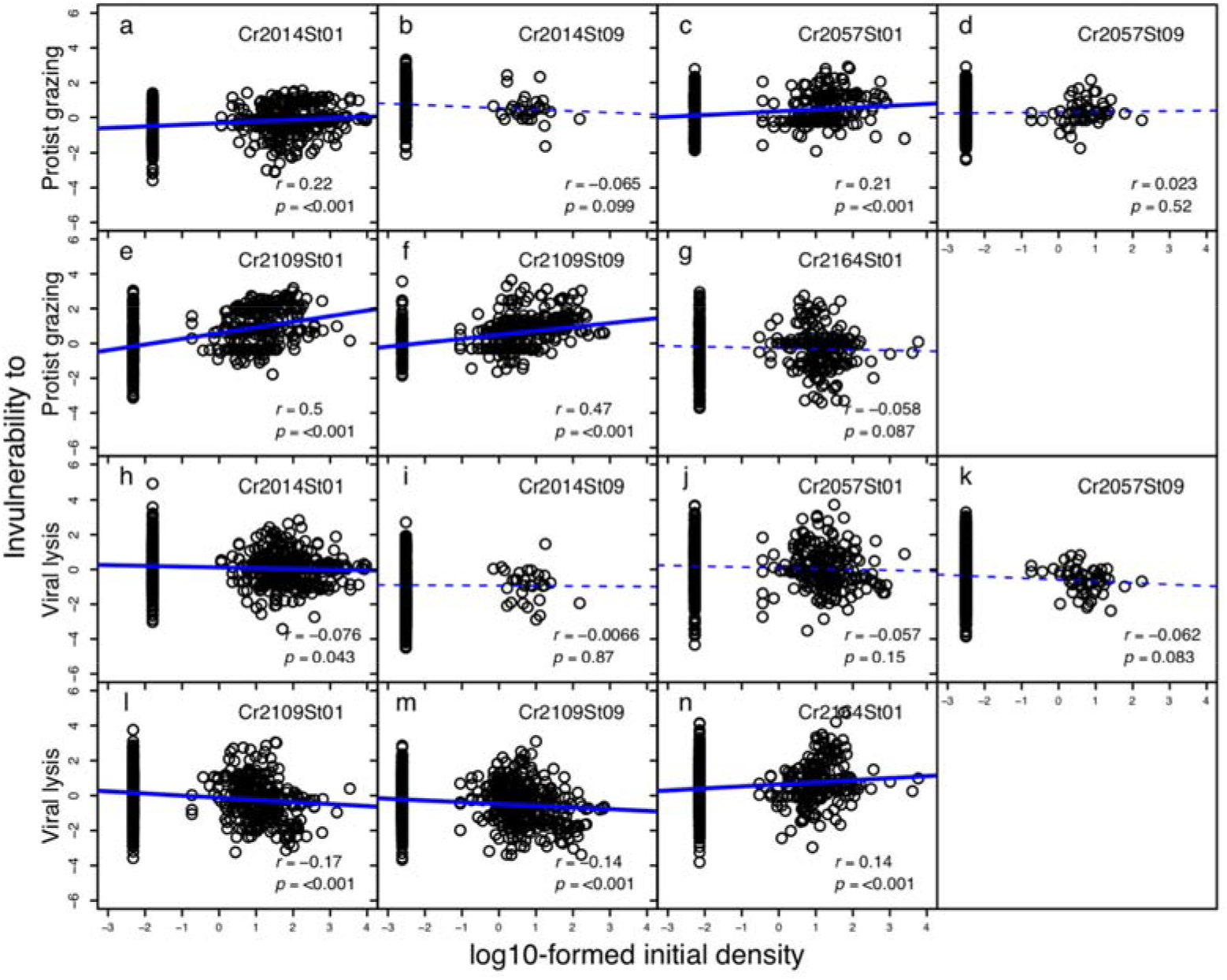
Relationships between log10-formed initial density and invulnerability to protist grazing or to viral lysis (hour^−1^), respectively, in each experiment. Values are the Pearson’s correlation coefficient (*r*) and its p value (*p*). Solid and dashed lines indicate significant (*p* < 0.05) and non-significant (*p* > 0.05) linear regression, respectively.

### Predation effect on bacterial diversity

Here, we explore how bacterial diversity responded to three types of predation pressures, respectively: (i) protists+viruses effect; (ii) protists effect; and (iii) viruses effect. The bacterial rank-abundance distribution (RAD) decay coefficient decreased (*LMM-p* <0.001) and evenness increased (*LMM-p* <0.001) when protists+viruses effect was increased. Similarly, when increasing only viruses effect, RAD decay coefficient decreased (*LMM-p* = 0.012) and evenness increased (*LMM-p* = 0.024). However, increasing only protists effect did not significantly change RAD decay coefficient and evenness (*LMM-p* = 0.335 and 0.237).

Bacterial richness increased when protists+viruses effect increased (*LMM-p* = 0.009) and when only protists effect was increased (*LMM-p* = 0.004). However, increasing only viruses effect did not change bacterial richness (*LMM-p* = 0.264). Therefore, Predictions 5 and 6 were only partially supported because protist grazing exhibited a more deterministic effect on bacterial richness whereas viral lysis had a major effect on bacterial RAD and evenness. When two predation pressures, i.e., both protist grazing and viral lysis, were combined, they increased bacterial diversity.

## Discussion

### The growth-based competitiveness-invulnerability trade-off in marine bacterial communities

Bacterial taxa with a higher predation-free growth rate had higher predation-caused mortality (Table 1 and Figure 1), indicating the presence of growth-based competitiveness-invulnerability trade-off in marine bacterial communities (supporting the Kill-the-Winner hypothesis). This finding was consistent with most studies inferring a possible competitiveness-invulnerability trade-off in marine microbes [22, 23, 24, 25, 26, 45]. Here, apparently for the first time, we obtained growth-based estimates on each bacterial taxon in field experiments to further explore the existence of competitiveness-invulnerability trade-offs in marine bacterial communities. Our findings provided empirical evidence supporting the Kill-the-Winner hypothesis that has been widely regarded as an important mechanism determining marine microbial assemblages and the performance of many ecosystem functions, but hitherto rarely verified in natural marine systems.

Furthermore, we inferred that viral lysis, and not protist grazing, was the major driver of the Kill-the-Winner mechanism in marine bacterial communities. This was supported by a strong and consistent growth-based competitive-invulnerability trade-off under viral lysis whereas this trade-off was weak or absent in some experiments when predation was contributed by protists grazing (Table 1 and Figure 1). Marine viruses are generally highly host-specific [21]. In empirical studies, virus-host interactions had a strong dependence on host growth rate [45]. Furthermore, there was a suggestion of higher taxonomic specificity in virus–bacteria relationships than protist–bacteria relationships [46]. Moreover, products released in the environment from viral lysis can promote growth of weak bacterial competitors due to increased nutrient availability [30, 47], which may further enhance the competitiveness-invulnerability trade-off. In contrast, marine protists are generally known to exploit a large spectrum of bacterial species [48, 49]. Therefore, a protist community may perform weak species selection on their prey when the proportion of specialist grazers in the community is low. In addition, specialist protists’ grazing can be size- or speed-selective [50, 51, 52] which may or may not be correlated to the bacterial predation-free growth rate examined in the present study. As such, contributions from “Kill-the-Winner” to the overall grazing selection on a bacterial community may be masked or weakened as a variety of selection occurs simultaneously.

### The density-based competitiveness-invulnerability trade-off in marine bacterial communities

Our findings did not support the existence of the competitiveness-invulnerability trade-off when taking bacterial initial density as a proxy to its competitiveness, regardless of whether predation was by protist grazing or viral lysis. This conclusion was inconsistent with prior studies that described viral infections being highly dependent on host density [25, 53], and species of protists that are specialized on the dominant bacterial species [54]. However, most previous studies were well-controlled laboratory experiments, whereas our results reflected the complexity of the natural system that would mask the density-based “Kill-the-Winner” in the present study.

Moreover, contrary to our predictions, the initially abundant bacterial species had relatively less predation-caused mortality in several experiments (Figure 2). Given this, we explained our findings from the perspective that full predation can best maintain bacterial community structure. That is, a weakened predation effect would result in decreased/increased relative abundance of initially abundant/rare bacterial taxa. This may be because our initial bacterial communities were collected from the field; therefore, the initial density of each bacterial taxon is the consequence after full predation selection. Consequently, dominant species could be both slower growing and more grazer/viruses defensive, as reported [24]. We thus acknowledge that our finding of no evidence to support density-based competitiveness-invulnerability trade-off may have been due to the limitations of our experimental settings; instead, our findings reflected that predation has a role in maintaining marine bacterial community structure. This was supported by the results where bacterial composition of the initial community was the closest to the composition incubated under a full predation effect (Figure S4).

### Predation effects on bacterial rank-abundance-distribution, evenness, and richness

When protist and virus density were increased together, there was a flattened bacterial rank-abundance distribution, as well as higher evenness and richness. These results were consistent with the expected consequences of the competitiveness-invulnerability trade-off in bacterial communities, where the predation effect can suppress growth of strong bacterial competitors that weakens the species exclusion effect from the competition. Although it is widely believed that predation is an important force shaping marine microbial diversity, whether it follows the often-cited Kill-the-Winner hypothesis remains unverified in natural systems. Here, our findings bridged the gap between the existence of competitiveness-invulnerability trade-off and predation-promoting effect on marine microbial diversity.

However, when protists were manipulated alone, there was no effect on bacterial RAD and evenness (Figure 3-4). One possible explanation may be that diluting protists alone could not be enough to change the strength of competitiveness-invulnerability trade-off and so as to affect bacterial RAD and evenness. Indeed, although in our study protist grazing generally preferred the fast-growing bacterial winner, this pattern was not consistent across experiments and the effect was relatively weak in comparison to viral lysis (Table 1 and Figure 1-2).

**Figure 3.**
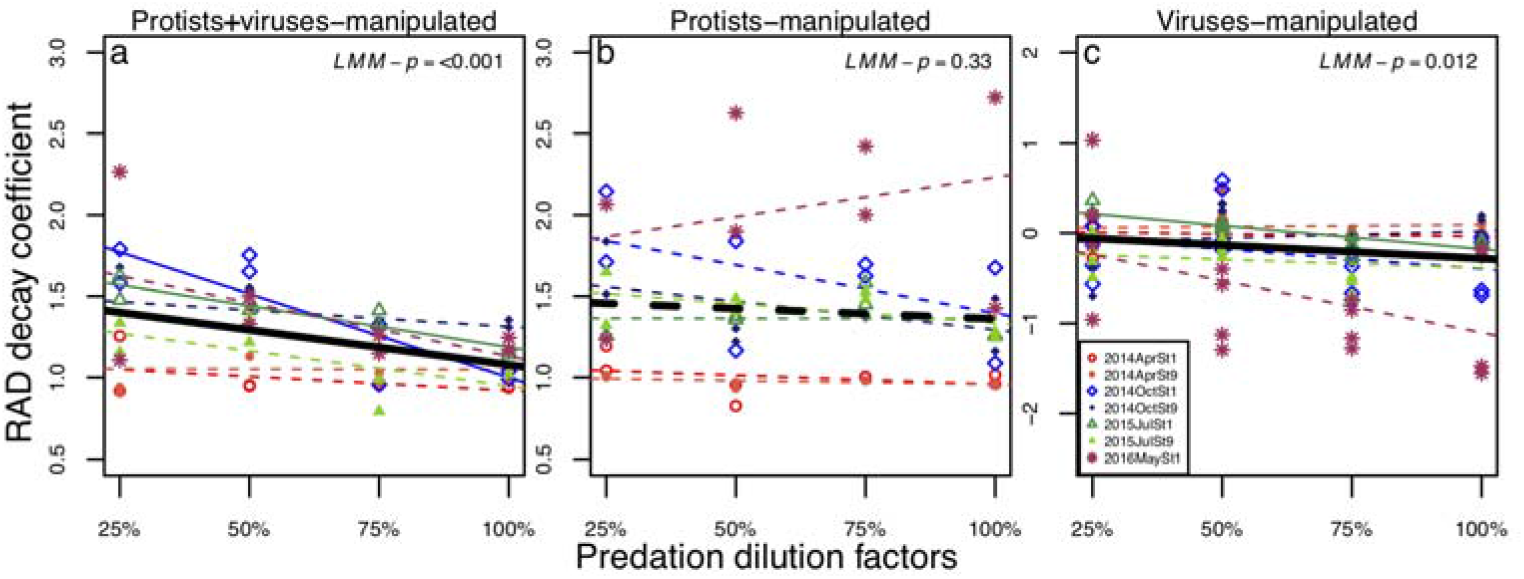
Relationships between predation dilution factors and bacterial RAD decay coefficient at T_12_ under (a) protists+viruses-manipulated, (b) protists-manipulated and, (c) viruses-manipulated treatments. Colors or symbols indicate different sets of experiments. Solid and dashed lines indicate significant (*p* < 0.05) and non-significant (*p* > 0.05) linear regression, respectively. Black bold solid and dashed lines indicate significant (*LMM-p* < 0.05) and non-significant (*LMM-p* > 0.05) linear regression estimated from linear-mixed effect model, with experiments as the random effect.

**Figure 4.**
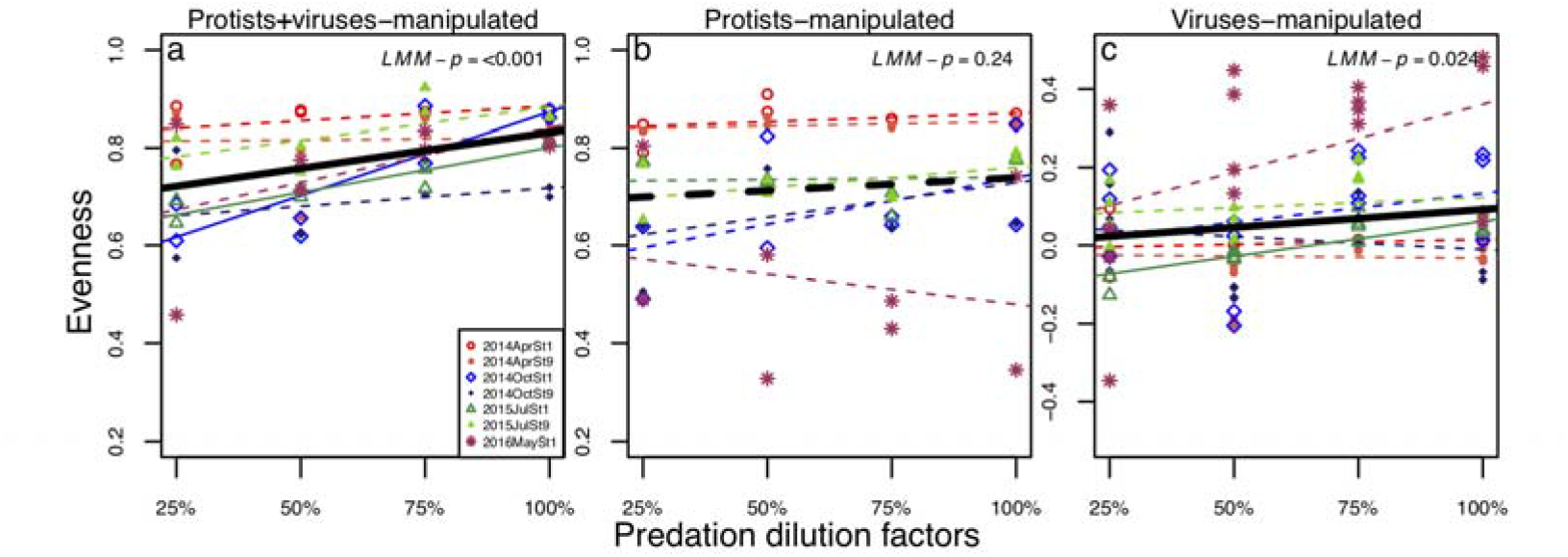
Relationships between predation dilution factors and bacterial evenness at T_12_ under (a) protists+viruses-manipulated, (b) protists-manipulated, and (c) viruses-manipulated treatments. Colors or symbols indicate different sets of experiments. Solid and dashed lines indicate significant (*p* < 0.05) and non-significant (*p* > 0.05) linear regression, respectively. Black bold solid and dashed lines indicate significant (*LMM-p* < 0.05) and non-significant (*LMM-p* > 0.05) linear regression estimated from linear-mixed effect model, with experiments as the random effect.

Nevertheless, bacterial richness increased with the increased protist grazing effect (Figure 5). Therefore, we speculated that protist grazing would maintain bacterial richness through other mechanisms rather than through its weak preference on fast-growing prey. For example, predation by generalist protists would suppress growth of overall bacterial biomass, allowing more remaining resources in the environment and supporting species co-existence.

**Figure 5.**
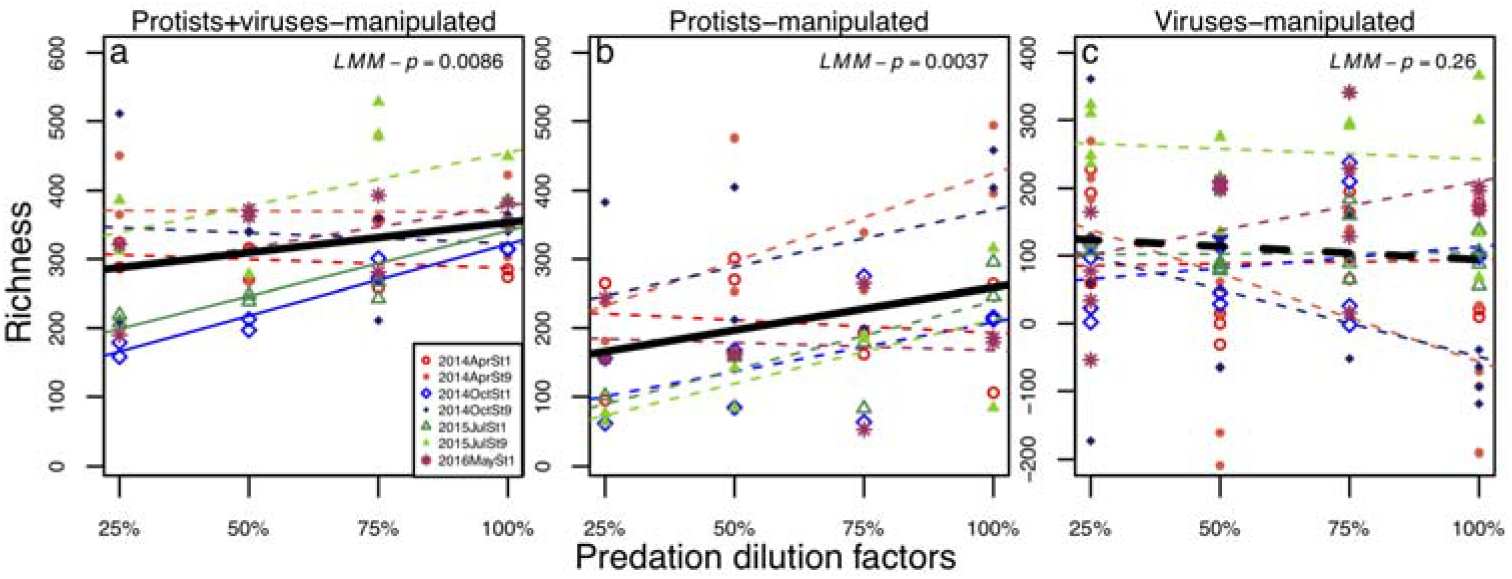
Relationships between predation dilution factors and bacterial richness at T_12_ under (a) protists+viruses-manipulated, (b) protists-manipulated, and (c) viruses-manipulated treatments. Colors or symbols indicate different sets of experiments. Solid and dashed lines indicate significant (*p* < 0.05) and non-significant (*p* > 0.05) linear regression, respectively. Black bold solid and dashed lines indicate significant (*LMM-p* < 0.05) and non-significant (*LMM-p* > 0.05) linear regression estimated from linear-mixed effect model, with experiments as the random effect.

On the contrary, when the viral effect increased, bacterial RAD and evenness became flatter and higher, respectively, whereas bacterial richness did not change. Therefore, we inferred that viral lysis had a critical role in controlling the dominance of the strong competitor; however, weakening the viral-driven competitiveness-invulnerability trade-off did not necessarily strengthen competition exclusion and lead to bacterial taxon loss.

We also acknowledge that we may have underestimated the viral lysis effects on bacterial diversity by subtracting the diversity indices of protists-manipulated treatment from protists+viruses-manipulated treatment. This concern was due to potential antagonistic interactions between protists and viruses, as suggested in previous studies, for instance, the competition on the same bacteria pools or intrigued predation where protists graze on free-living virus and viral-infected bacteria [55, 56, 57]. This would also explain why many experiments exhibited no significant viral lysis effect on bacterial diversity (Figure 3-5). Experiments manipulating only viral lysis effect would provide better evidence [23, 24], but are technically challenging.

In addition, our seven experiments were conducted in different sites or seasons, and in many, there was no predation effect on bacterial diversity (Figure 3-5). Perhaps predation effects were weakened or masked by other environmental factors [7, 8, 9]. Furthermore, we incubated marine microbes under *in-situ* temperature and in a closed system, without nutrient and particle exchange. Low temperatures, depleted nutrients, and accumulated metabolic waste can occur and weaken bacterial productivity to exhibit a significant growth response to predation. Yet, there was no apparent relationship between environment nutrient concentration, bacterial initial abundance, and richness versus how predation affects bacterial diversity (Table S2), perhaps, owing to our limited number of experimental sets.

Finally, we also acknowledge that a potential “bottle effect” may have existed in our experiments. The bottle effect can result in a bacterial community change due to an incubation environment in the bottles that may weaken or mask bacterial community responses to the manipulated predation effect, as suggested [58, 59]. To exam a potential bottle effect in the present experiment, we investigated the ranking of ASVs (ranked by relative abundance) in T_0_ versus T_12_ when without predation manipulation (under 100% predation dilution factor, Figure S5). The ASVs with higher ranking at T_0_ generally had a higher ranking in T_12_ when without predation manipulation, indicating that, generally, ASVs ranking was not dramatically influenced by the incubation environment (Figure S5). Nevertheless, some ASVs did have a large ranking shift, potentially caused by a bottle effect. For a better quantitative examination, we analyzed the degree of ranking change for each ASV by calculating | T_0-Rank_/T_0-MaxRank_−T_12-100%-Rank_/T_12-100%-MaxRank_| for each ASV; where T_0-Rank_ and T_12-100%-Rank_ are the ranking at T_0_ and T_12_; and T_0-MaxRank_ and T_12-100%-MaxRank_ are the maximum ranking at T_0_ and at T_12_, respectively. When consider all ASVs, only 9.9% of ASVs had < 10% ranking change; nevertheless, in the top-100 rank ASVs (based on relative abundance across all experiments) which contained 86% of total reads, 87 ASVs had <10% of ranking change. Therefore, we concluded that bottle effects potentially existed and influenced bacterial community components; however, bottle effects did not alter the major component of most bacterial abundance in our study. In addition, even if a bottle effect caused a large ranking shift in rare bacterial taxa, it did not mask the overall bacterial community responses to the manipulated predation effects.

### Predation effects on bacterial composition

A final check was conducted to examine if the Bray-Curtis dissimilarity between bacterial communities at T_12_ and T_0_ varied under different predation dilution factors. The Bray-Curtis dissimilarities decreased with predation, regardless of protists-manipulated or protists+viruses-manipulated treatments in LMM analysis (Figure S4). This finding agreed with most studies, suggesting that predation maintained bacterial composition [15, 60].

## Materials and methods

### Study sites and predation manipulation experiments

In total, seven sets of predation manipulation experiments were conducted in April 2014, October and July 2015, and May 2016 at two stations in the East China Sea (Figure S1). Bacterial communities were collected at the surface layer (3 to 5 m) by Go-Flo bottles mounted on a CTD-equipped rosette (Sea-Bird Electronics, Bellevue, WA, USA). For each set of predation manipulation experiments, three types of seawater samples were prepared: (i) macro-plankton-free (20-μm filtered); (ii) protists- and bacteria-free (0.22-μm filtered); and (iii) particle-free seawater (30-kDa filtered). These three types of seawater were used for creating two combinations of predation manipulation treatments: (i) Protists-manipulated (macro-plankton-free seawater diluted by grazing- and bacteria-free water) and (ii) Protists+viruses-manipulated (macro-plankton-free seawater diluted by particle-free seawater). This predation manipulation technique followed previous studies that assumed filtration and manipulation processes do not affect dissolved-organic matter and bacterial growth, but decrease bacterial mortality caused by protists grazing and viral lysis due to decreased encounter rates between bacteria and predators [23, 31, 32, 33]. For each combination of predation manipulation, a 4-point dilution series was generated, with 25, 50, 75 and 100% of the original predation effect remaining (predation dilution factors) (Figure S2). The mixtures were incubated for 12 h in duplicate in sterilized 500-ml polycarbonate bottles at *in situ* temperature in a cabin, with water flow taken on board from the sea surface layer and under natural light. For bacterial enumeration samples, 2 ml of seawater from each 500-ml bottle were collected at the beginning (T_0_) and after 12-h incubation (T_12_) and fixed by paraformaldehyde solution with a final concentration 0.2% [34]. Therefore, 32 samples were collected from each experiment, which contains two predation manipulation treatments (protists-manipulated and viruses-manipulated treatments) with four predation dilution factors in duplicates collected at T_0_ and T_12_. In total, 224 samples were collected in seven experiments.

For 16S rDNA sequencing samples, ~ 10 L of macro-plankton-free seawater (20-μm filtered) representing the community composition in each experiment at T_0_, and the rest of the seawater in each of the 16 half-litter bottles (two predation manipulation treatments with four predation dilution factors in duplicates) at T_12_ (representing the community composition after 12-h incubation), were filtered through 0.2-μm pore size polycarbonate membranes, respectively. Consequently, 17 community samples were collected from each experiment; thus, a total of 119 community samples were collected from seven experiments. All seawater and filter samples were frozen with liquid nitrogen onboard and stored at −80°C until the next process.

### Bacteria enumeration, DNA extraction, PCR, sequencing and bioinformatics

To enumerate bacterial, 2 ml of paraformaldehyde-fixed seawater samples was stained with SYBR green (Molecular Probes Inc., USA) for 15 min in the dark and enumerated with a FACSAria flow cytometer (Becton Dickinson, USA).

For 16S rRNA gene sequencing, total DNA was extracted from the 0.2-μm pore size membranes using DNeasy PowerWater Kit (Qiagen, Hilden, Germany) according to manufacturer’s instructions. For generating libraries, DNA extracts from the membranes were used as templates of polymerase chain reaction (PCR) to amplify the V5–V6 region [35] of 16S rDNA, followed by a second PCR to generate amplicons with unique dual-index for each sample. However, we failed to amplify one 16S rDNA sample (the 25% predation dilution factor under the protists-manipulated treatment in the experiment 2015JulSt1) in the first PCR; thus, only 118 samples proceeded to the next step. PCR purification was done after the first and second PCRs using Agencourt AMPure XP beads (Beckman Coulter, Indianapolis, Indiana). The purified DNA samples were pooled with approximately equal DNA quantity measured with Qubit dsDNA BR Assay Kit (Thermo Fisher Scientific, Waltham, Massachusetts; see Supplementary Information for details). Sequencing was carried out using the Illumina MiSeq platform, producing 2 × 300 bp paired-end reads. Raw sequence data were deposited in the NCBI Sequence Read Archive (SRA) under accession number PRJNA749329.

Raw sequences were processed using the Quantitative Insights Into Microbial Ecology 2 (QIIME 2 v2018.8) pipeline [36], following a protocol that processed with DADA2 with standard parameters [37] for removing primers, reads trimming (forward and reverse reads were truncated reads at positions 210 and 140 where quality score start to crush below 25, respectively), errors correction, merging read pairs, removing possible PCR chimeras (consensus method), and generating amplicon sequence variant (ASV) representative sequences and tables. To identify and remove potential mitochondria and chloroplast ASV, representative sequences were classified with a Naive Bayes classifier [38] fitted with 16S rRNA gene sequences extracted from SILVA v132 database [39] based on the PCR primers used. Singleton ASVs (which may be created when merging paired-end sequences) across all experiments were removed.

### ASV table rarefaction and relative abundance estimates for the competitiveness-invulnerability trade-off analysis

The singleton-free ASV table was separated into seven sets of experiments and rarefied based on the lowest number of reads among all communities in each experiment (Table S1). If the relative abundance of the bacterial taxon (each ASV) in a community was 0, we replaced it with the relative abundance using the ASV table prior to rarefaction in that community. Furthermore, if the relative abundance remained 0 in the ASV table prior to rarefaction, we set it as 10^−8^. This value was determined according to the findings that the smallest relative abundance before rarefaction for each of the seven sets of experiments was ~ 10^−7^ (Table S2). A sensitivity test on setting the zero relative abundance in both rarefied and non-rarefied ASV table as 10^−7^ was also conducted; results indicated a consistent conclusion on the competitiveness-invulnerability relationship (Table S2). This adjustment on zero relative abundance is necessary to alleviate impacts from sampling effects that would otherwise neglect responses of bacterial taxa susceptible to predation manipulation, but extremely rare. For example, potential predation-susceptible taxa might be initially too rare to be detected, but subsequently appear after 12-h incubation under diluted predation treatments (i.e., 25, 50 and 75% diluted treatments, ranging from 56.9 to 80.9% of total ASVs with 5.3 to 19% of total reads across experiments). Alternatively, the bacterial taxa existed initially but was not detected after 12 h under a diluted predation effect (<0.6% of ASVs with < 0.001% of total reads across all experiments, Table S3), i.e., the taxa are potentially predation-defenders but susceptible to competition.

### Estimates of competitiveness and invulnerability to predation of each bacterial ASV

***The initial density*** of each bacterial ASV was calculated by multiplying its relative abundance (estimated from 16s rDNA sequencing) and the mean of the density of the whole community (enumerated with flow cytometry) at T_0_. The relationship between ***per-capita net growth rate*** versus ***predation dilution factors (PNGR-PDF)*** were analyzed for each bacterial ASV (hereafter, the PNGR-PDF relationship) for both predation manipulation treatments (protists-manipulated and protists+viruses-manipulated) (Figure S3). The ***per-capita net growth rate (r)*** was estimated using bacterial population size initially (N_0_) and after 12-h incubation (N_12_) where ***r*** = ln(N_12_/N_0_)/(*12 h*). This calculation assumes ***r*** does not change with population size with equation: N_t_ = N_0_\**re^t^*, where N_t_ is the population size at time *t* [40, 41]. Bacterial population size was estimated by multiplying the relative abundance and the absolute abundance of the whole community. Two linear regressions of the ***PNGR-PDF relationship*** for two predation manipulation treatments (with two replicates for each) were analyzed. ***Bacterial predation-free growth rate*** was estimated as the *y*-intercept of the PNGR-PDF relationship under the protists+viruses-manipulated treatment. ***Bacterial invulnerability to protist grazing*** was estimated by the regression slope of PNGR-PDF relationship under the protists-manipulated treatment, indicating the degree of bacterial growth rate in response to increased protist grazing. Finally, ***bacterial invulnerability to viral lysis*** was calculated by subtracting the protists-manipulated PNGR-PDF regression slope from the protists+viruses-manipulated PNGR-PDF regression slope [31, 33], representing the degree of bacterial growth rate in response to the increased viral lysis effect.

### Bacterial community indices

Here, we investigated three bacterial diversity indices: the decay coefficient of species rank abundance distribution (RAD), evenness and richness. The decay coefficient of RAD and evenness were estimated based on the rank-normalized RAD, through which the single-free ASV tables were scaled by taking the average of 1000 times subsampling with the lowest number of RAD rank (lowest number of species among communities) at T_12_ in each experiment and then normalized between 0 and 1 (Table S1) using “*RADanalysis*” package in R program. This allowed a quantitative comparison of RAD structures and evenness among communities with controlled richness [42]. To describe the RAD structure, the RAD decay coefficient was represented by the estimated decay coefficient obtained from fitting the Zipf model to the normalized RADs using the “*radfit*” package in the R program. A community with a higher RAD decay coefficient indicates that the frequency of species decreases dramatically along with rank (a steeper RAD) whereas a lower RAD decay coefficient represents a smaller difference among species frequency (a flatter RAD). Evenness was estimated using the “*vegan*” package in R program with rank-normalized RAD. Bacterial richness was estimated from the singleton-free ASV tables rarefied at the lowest number of reads among communities at T_12_ in each experiment (Table S1). Bacterial richness was the number of ASV of each community.

Finally, Bray–Curtis dissimilarity between communities collected at T_0_ and T_12_ was calculated to check predation effects on bacterial species composition. This analysis was done based on the singleton-free ASV tables that were rarefied at the lowest number of reads among all communities in each experiment (Table S1). Bray– Curtis dissimilarities were calculated based on the relative abundance of various ASVs using “*vegan*” package in the R program.

### Data analyses

Hypothesis I aimed at investigating the bacterial competitiveness-invulnerability trade-off. For each experiment, we determined linear regression and the Pearson’ correlation coefficient between bacterial competitiveness (as independent variable, for predation-free growth rate and initial abundance, respectively) versus invulnerability to predation (as dependent variable, for mortality caused by protists grazing and viral lysis, respectively). Then, pooling all these relationships across communities, we analyzed competitiveness-invulnerability relationships with a linear mixed-effects model (LMM) considering random intercepts across seven experiments using “*nlme*” package in R, with bacterial competitiveness as independent variable, invulnerability to predation as dependent variable, and experiments as the random effect. Additionally, to detect a potential artifactual slope-intercept relationship when analyzing competitiveness-invulnerability trade-off, a permutation test was conducted. This additional analysis was due to the concern that the estimates of slope (invulnerability to predation) and intercept (predation-free growth rate) were from two closely related regression lines. Specifically, for each predation effect (protists grazing or viral lysis, respectively), a null model was generated using the linear regression estimates with randomly shuffled dependent variables (the invulnerability to predation) across dilution factors while independent variables (predation-free growth rate) remain fixed and was repeated 1000 times. Observed values were statistically tested if significantly different from the mean of null models with Z-score and its p-value.

In Hypothesis II, we analyzed the relationship between predation dilution factors versus the three bacterial diversity indices (RAD decay coefficient, evenness, or richness) after 12-h incubation under three combinations of predation effect (protists+viruses, protists or viruses), respectively. These relationships were analyzed with LMM analysis as well, with predation dilution factors as independent variables, diversity indices as dependent variables, and experiments as the random effect. Three types of predation pressures on bacterial diversity were analyzed: (i) protists+viruses effect; (ii) protists effect; and (iii) viruses effect. Effects from protists and protists+virus were examined from protists-manipulated and protists+viruses-manipulated treatments, respectively. Bacterial diversity estimates under viral lysis effect were obtained by subtracting each diversity estimate in protists-manipulated treatment from each diversity estimate under protists+viruses-manipulated treatment. There are two replicates in each treatment, resulting in four diversity estimates under each viruses-manipulated dilution factors. Finally, we investigated how bacterial composition similarity between initial and 12-h incubation (Bray-Curtis dissimilarity) varied under the four degrees of predation dilution factors, with protists-manipulated and protists+viruses-manipulated treatments. Details of the data analysis and R scripts are provided online (https://github.com/jinnyyang/Marine-Bacteria-C-I-trade-off).

### Environmental variables

Temperature was recorded by the CTD profiler. Nitrite and nitrate concentrations were measured by the pink dye method, and phosphate concentrations were measured by molybdenum blue, using standard methods [43]. Environmental conditions during the experiments are summarized in Table S4.

## Acknowledgements

We thank Ching-Wei Hsu for assistance to conduct experiments and sampling, Hon-Tsen Yu for providing facilities and advice on laboratory work, and Sara Jackrel and Vincent Denef for comments. This work was supported by the National Center for Theoretical Sciences, Foundation for the Advancement of Outstanding Scholarship, and the Ministry of Science and Technology, Taiwan.

## References

1. Prosser JI, Bohannan BJ, Curtis TP, Ellis RJ, Firestone MK, Freckleton RP, Green JL, Green LE, Killham K, Osborn AM, Solan M, van der Gast CJ, Young, JPW. The role of ecological theory in microbial ecology. Nature Reviews Microbiology. 2007;5:384–92. dio:10.1038/nrmicro1643.

2. Hutchins DA, Fu F. Microorganisms and ocean global change. Nature Microbiology. 2017;2:1–11. doi:10.1038/nmicrobiol.2017.58.

3. Cavicchioli R, Ripple WJ, Timmis KN, Azam F, Bakken LR, Baylis M, Behrenfeld MJ, Boetius A, Boyd PW, Classen AT, Crowther TC, Danovaro R, Foreman CM, Huisman J, Hutchins DA, Jansson JK, Karl DM, Koskella B, Welch DM, Martiny JB, Moran MA, Orphan VJ, Reay DS, Remais JV, Rich VI, Singh BK, Stein LY, Stewart FJ, Matthew B. Sullivan MB, van Oppen MJ, Scott C. Weaver SC, Webb EA, Webster NS. Scientists’ warning to humanity: microorganisms and climate change. Nature Reviews Microbiology. 2019;17:569–86. doi:10.1038/s41579-019-0222-5.

4. Azam F, Fenchel T, Field JG, Gray J, Meyer-Reil L, Thingstad F. The ecological role of water-column microbes in the sea. Marine Ecology Progress Series. 1983;20:257–63. doi:10.3354/meps010257.

5. Hagström A, Azam F, Andersson A, Wikner J, Rassoulzadegan F. Microbial loop in an oligotrophic pelagic marine ecosystem: possible roles of cyanobacteria and nanoflagellates in the organic fluxes. Marine ecology progress series. Oldendorf. 1988;49:171–78. doi:10.3354/meps049171.

6. Fuhrman JA, Cram JA, Needham DM. Marine microbial community dynamics and their ecological interpretation. Nature Reviews Microbiology. 2015;13:133–46. doi:10.1038/nrmicro3417.

7. Nogales B, Lanfranconi MP, Piña-Villalonga JM, Bosch R. Anthropogenic perturbations in marine microbial communities. FEMS Microbiology Reviews. 2011;35:275–98. doi:10.1111/j.1574-6976.2010.00248.x.

8. Yeh YC, Peres-Neto PR, Huang SW, Lai YC, Tu CY, Shiah FK, Gong GC, Hsieh CH. Determinism of bacterial metacommunity dynamics in the southern East China Sea varies depending on hydrography. Ecography. 2015;38:198–212. doi:10.1111/ecog.00986.

9. York A. Marine microbial diversity from pole to pole. Nature Reviews Microbiology. 2020;18:3. doi:10.1038/s41579-019-0304-4.

10. Fuhrman JA. Microbial community structure and its functional implications. Nature. 2009;459:193–99. doi:10.1038/nature08058.

11. Ratzke C, Barrere J, Gore J. Strength of species interactions determines biodiversity and stability in microbial communities. Nature Ecology & Evolution. 2020;4:376–383. doi:10.1038/s41559-020-1099-4.

12. Chang FH, Yang JW, Liu AC, Lu HP, Gong GC, Shiah FK, Hsieh CH. Community assembly processes as a mechanistic explanation of the predator-prey diversity relationship in marine microbes. Frontiers in Marine Science. 2021;8:601. doi:10.3389/fmars.2021.651565.

13. Kneitel JM, Chase JM. Trade-offs in community ecology: linking spatial scales and species coexistence. Ecology Letters. 2004;7:69–80. doi:10.1046/j.1461-0248.2003.00551.x.

14. Litchman E, Klausmeier CA, Schofield OM, Falkowski PG. The role of functional traits and trade-offs in structuring phytoplankton communities: scaling from cellular to ecosystem level. Ecology Letters. 2007;10:1170–81. doi:10.1111/j.1461-0248.2007.01117.x.

15. Winter C, Bouvier T, bauer MG, Thingstad TF. Trade-offs between competition and defense specialists among unicellular planktonic organisms: the “killing the winner” hypothesis revisited. Microbiology and Molecular Biology Reviews. 2010;74:42–57. doi:10.1128/MMBR.00034-09.

16. Thingstad TF. Elements of a theory for the mechanisms controlling abundance, diversity, and biogeochemical role of lytic bacterial viruses in aquatic systems. Limnology and Oceanography. 2000;45:1320–28. doi:10.4319/lo.2000.45.6.1320.

17. Edwards KF, Klausmeier CA, Litchman E. A three-way trade-off maintains functional diversity under variable resource supply. The American Naturalist. 2013;182:786–800. doi:10.1086/673532.

18. Litchman E, Edwards KF, Klausmeier CA. Microbial resource utilization traits and trade-offs: implications for community structure, functioning, and biogeochemical impacts at present and in the future. Frontiers in Microbiology. 2015;6:254. doi:10.3389/fmicb.2015.00254.

19. Lenski RE. Experimental studies of pleiotropy and epistasis in Escherichia coli. Variation in competitive fitness among mutants resistant to virus T4. Evolution. 1988;42:425–32. doi:10.1111/j.1558-5646.1988.tb04149.x.

20. Jessup CM, Bohannan BJ. The shape of an ecological trade-off varies with environment. Ecology Letters. 2008;11:947–59. doi:10.1111/j.1461-0248.2008.01205.x.

21. Fuhrman J, Schwalbach M. Viral influence on aquatic bacterial communities. The Biological Bulletin. 2003;204:192–5. doi:10.2307/1543557.

22. Zhang R, Weinbauer MG, Qian PY. Viruses and flagellates sustain apparent richness and reduce biomass accumulation of bacterioplankton in coastal marine waters. Environmental Microbiology. 2007;9:3008–18. doi:10.1111/j.1462-2920.2007.01410.x.

23. Bonilla-Findji O, Herndl GJ, Gattuso JP, Weinbauer MG. Viral and flagellate control of prokaryotic production and community structure in off-shore Mediterranean waters. Applied and Environmental Microbiology. 2009;75:4801–12. doi:10.1128/AEM.01376-08.

24. Cram JA, Parada AE, Fuhrman JA. Dilution reveals how viral lysis and grazing shape microbial communities. Limnology and Oceanography. 2016;61:889–905. doi:10.1002/lno.10259.

25. Fuhrman JA, Noble RT. Causative agents of bacterial mortality and the consequences to marine food webs. Microbial Biosystems: New Frontiers. 2000:145–51.

26. Suttle CA. Marine viruses—major players in the global ecosystem. Nature Reviews Microbiology. 2007;5:801–12. doi:10.1038/nrmicro1750.

27. Bouvier T, Del Giorgio PA. Key role of selective viral-induced mortality in determining marine bacterial community composition. Environmental Microbiology. 2007;9:287–97. doi:10.1111/j.1462-2920.2006.01137.x.

28. Unrein F, Massana R, Alonso-Sáez L, Gasol JM. Significant year-round effect of small mixotrophic flagellates on bacterioplankton in an oligotrophic coastal system. Limnology and Oceanography. 2007;52:456–69. doi:10.4319/LO.2007.52.1.0456.

29. Schmoker C, Hernández-León S, Calbet A. Microzooplankton grazing in the oceans: impacts, data variability, knowledge gaps and future directions. Journal of Plankton Research. 2013;35:691–706. doi:10.1093/plankt/fbt023

30. Zhang R, Li Y, Yan W, Wang Y, Cai L, Luo T, Li H, Weinbauer MG, Jiao N. Viral control of biomass and diversity of bacterioplankton in the deep sea. Communications Biology. 2020;3:1–10. doi:10.1038/s42003-020-0974-5.

31. Evans C, Archer SD, Jacquet S, Wilson WH. Direct estimates of the contribution of viral lysis and microzooplankton grazing to the decline of a Micromonas spp. population. Aquatic Microbial Ecology. 2003;3:207–19. doi:10.3354/ame030207.

32. Kimmance SA, Wilson WH, Archer SD. Modified dilution technique to estimate viral versus grazing mortality of phytoplankton: limitations associated with method sensitivity in natural waters. Aquatic Microbial Ecology. 2007;49:207–22. doi:10.3354/ame01136.

33. Tsai AY, Gong GC, Hung J. Seasonal variations of virus- and nanoflagellate-mediated mortality of heterotrophic bacteria in the coastal ecosystem of subtropical western Pacific. Biogeosciences. 2013;10:3055–65. doi:10.5194/bg-10-3055-2013.

34. Chung CC, Gong GC, Huang CY, Lin JY, Lin YC. Changes in the Synechococcus assemblage composition at the surface of the East China Sea due to flooding of the Changjiang river. Microbial Ecology. 2015;70:677–88. doi:10.1007/s00248-015-0608-5.

35. Cai L, Ye L, Tong AHY, Lok S, Zhang T. Biased diversity metrics revealed by bacterial 16S pyrotags derived from different primer sets. PloS One. 2013;8:e53649. doi:10.1371/journal.pone.0053649.

36. Bolyen E, Rideout JR, Dillon MR, Bokulich NA, Abnet CC, Al-Ghalith GA, et al. Reproducible, interactive, scalable and extensible microbiome data science using QIIME 2. Nature Biotechnology. 2019;37:852–57. doi:10.1038/s41587-019-0209-9.

37. Callahan BJ, McMurdie PJ, Rosen MJ, Han AW, Johnson AJA, Holmes SP. DADA2: high-resolution sample inference from Illumina amplicon data. Nature Methods. 2016;13:581–83. doi:10.1038/nmeth.3869.

38. Pedregosa F, Varoquaux G, Gramfort A, Michel V, Thirion B, Grisel O, et al. Scikit-learn: Machine learning in Python. The Journal of Machine Learning Research. 2011;12:2825–2830.

39. Quast C, Pruesse E, Yilmaz P, Gerken J, Schweer T, Yarza P, Peplies J, Glöckner FO. The SILVA ribosomal RNA gene database project: improved data processing and web-based tools. Nucleic Acids Research. 2012;41:D590–6. doi:10.1093/nar/gks1219.

40. Landry M, Hassett R. Estimating the grazing impact of marine micro-zooplankton. Marine Biology. 1982;67:283–88. doi:10.1007/BF00397668.

41. Salat J, Marrase C. Exponential and linear estimations of grazing on bacteria: effects of changes in the proportion of marked cells. Marine Ecology Progress Series. 1994;104:205. doi: 10.3354/meps104205.

42. Saeedghalati M, Farahpour F, Budeus B, Lange A, Westendorf AM, Seifert M, Küppers R, Hoffmann D. Quantitative comparison of abundance structures of generalized communities: from B-cell receptor repertoires to microbiomes. PLoS Computational Biology. 2017;13:e1005362. doi:10.1371/journal.pcbi.1005362.

43. Gong GC, Wen YH, Wang BW, Liu GJ. Seasonal variation of chlorophyll a concentration, primary production and environmental conditions in the subtropical East China Sea. Deep Sea Research Part II: Topical Studies in Oceanography. 2003;50:1219–36.doi:10.1016/S0967-0645(03)00019-5.

44. Storesund JE, Erga SR, Ray JL, Thingstad TF, Sandaa RA. Top-down and bottom-up control on bacterial diversity in a western Norwegian deep-silled fjord. FEMS Microbiology Ecology. 2015;91:fiv076. doi:10.1093/femsec/fiv076.

45. Middelboe M. Bacterial growth rate and marine virus–host dynamics. Microbial Ecology. 2000;40:114–24. doi:10.1007/s002480000050.

46. Chow CET, Kim DY, Sachdeva R, Caron DA, Fuhrman JA. Top-down controls on bacterial community structure: microbial network analysis of bacteria, T4-like viruses and protists. The ISME journal. 2014;8:816–29. doi:10.1038/ismej.20.

47. Coutinho FH, Silveira CB, Gregoracci GB, Thompson CC, Edwards RA, Brussaard CP, Dutilh BE, Thompson FL. Marine viruses discovered via metagenomics shed light on viral strategies throughout the oceans. Nature Communications. 2017;8:1–12. doi:10.1038/ncomms15955.

48. Johnke J, Baron M, de Leeuw M, Kushmaro A, Jurkevitch E, Harms H, Chatzinotas A. A generalist protist predator enables coexistence in multi-trophic predator-prey systems containing a phage and the bacterial predator bdellovibrio. Frontiers in Ecology and Evolution. 2017;5:124. doi:10.3389/fevo.2017.00124.

49. Bjorbækmo MFM, Evenstad A, Røsæg LL, Krabberød AK, Logares R. The planktonic protist interactome: where do we stand after a century of research? The ISME journal. 2020;14(2):544–59. doi:10.1038/s41396-019-0542-5.

50. Güde H. The role of grazing on bacteria in plankton succession. Plankton ecology. Springer; Heidelberg, Berlin, Germany: 1989. pp. 337–364.

51. Jürgens K, Matz C. Predation as a shaping force for the phenotypic and genotypic composition of planktonic bacteria. Antonie Van Leeuwenhoek. 2002;81:413–34. doi:10.1023/a:1020505204959.

52. Batani G, Pérez G, Martínez de la Escalera G, Piccini C, Fazi S. Competition and protist predation are important regulators of riverine bacterial community composition and size distribution. Journal of Freshwater Ecology. 2016;31:609–23. doi:10.1080/02705060.2016.1209443.

53. Maslov S, Sneppen K. Population cycles and species diversity in dynamic Kill-the-Winner model of microbial ecosystems. Scientific Reports. 2017;7:1–8. doi:10.1038/srep39642.

54. Bell T, Bonsall MB, Buckling A, Whiteley AS, Goodall T, Griffiths RI. Protists have divergent effects on bacterial diversity along a productivity gradient. Biology Letters. 2010;6:639–42. doi:10.1098/rsbl.2010.0027.

55. Miki T, Yamamura N. Intraguild predation reduces bacterial species richness and loosens the viral loop in aquatic systems:‘kill the killer of the winner’hypothesis. Aquatic Microbial Ecology. 2005;40:1–12. doi:10.3354/AME040001.

56. Deng L, Krauss S, Feichtmayer J, Hofmann R, Arndt H, Griebler C. Grazing of heterotrophic flagellates on viruses is driven by feeding behaviour. Environmental Microbiology Reports. 2014;6:325–30. doi:10.1111/1758-2229.12119.

57. Bouvy M, Bettarel Y, Bouvier C, Domaizon I, Jacquet S, Le Floc’h E, Montaié H, Mostajir B, Sime-Ngando T, Torréton JP, Vidussi F, Bouvier T. Trophic interactions between viruses, bacteria and nanoflagellates under various nutrient conditions and simulated climate change. Environmental Microbiology. 2011;13:1842–57. doi:10.1111/j.1462-2920.2011.02498.x.

58. Agis M, Granda A, Dolan JR. A cautionary note: examples of possible microbial community dynamics in dilution grazing experiments. Journal of Experimental Marine Biology and Ecology. 2007;341:176–83. doi:10.1016/j.jembe.2006.09.002.

59. Hammes F, Vital M, Egli T. Critical evaluation of the volumetric “bottle effect” on microbial batch growth. Applied and Environmental Microbiology. 2010;76:1278–81. doi:10.1128/AEM.01914-09.

60. Schwalbach MS, Hewson I, Fuhrman JA. Viral effects on bacterial community composition in marine plankton microcosms. Aquatic Microbial Ecology. 2004;34:117–27. doi:10.3354/AME034117.

