## Supplementary information for "Trade-off between competition ability and invulnerability to predation in marine microbes; protist grazing versus viral lysis effects"

**Supplementary figures and tables**

Table S1. Rarefied number of reads and fixed RAD rank for ASV table standardization for each experiment. Note that: (a) bacterial net growth rates were estimated from communities rarefied with the lowest number of reads among all samples in each experiment; (b) bacterial richness was estimated from communities rarefied with the lowest number of reads among samples at $T_{12}$ in each experiment; and (c) the RAD decay coefficient and evenness were estimated based on communities scaled by the lowest number of RAD rank (number of species) among samples at $T_{12}$ in each experiment.

| **Experiment** | (a) Rarefied number of reads for net growth rate estimates | (b) Rarefied number of reads for richness at $T_{12}$ estimates | (c) Fixed RAD rank for RAD decay coefficient and evenness at $T_{12}$ estimates |
|---|---|---|---|
| 2014AprSt1 | 6751 | 6751 | 96 |
| 2014AprSt9 | 1598 | 17502 | 182 |
| 2014OctSt1 | 7140 | 7140 | 61 |
| 2014OctSt9 | 3188 | 16560 | 151 |
| 2015JulSt1 | 12355 | 12355 | 85 |
| 2015JulSt9 | 6467 | 6467 | 66 |
| 2016MaySt1 | 13343 | 13343 | 54 |

Table S2. The smallest value of relative abundance calculated from the non-rarefied ASV table for each set of experiments, as well as the growth-based competitiveness-invulnerability relationship when setting zero relative abundance as $10^{-7}$. The smallest relative abundance for each experiment was calculated by the sum of the reads of each bacterial taxon (ASV) across all communities, divided by the total number of reads in each experiment. The growth-based competitiveness-invulnerability relationships are represented by the Pearson's correlation coefficient (Cor.) and its *p* value.

| | Smallest relative abundance (non-rarefied) | Growth-based competitiveness-invulnerability relationship | | | | Density-based competitiveness-invulnerability relationship | | | |
|---|---|---|---|---|---|---|---|---|---|
| | | Under protists grazing | | Under viral lysis | | Under protists grazing | | Under viral lysis | |
| | | Cor. | *p* value | Cor. | *p* value | Cor. | *p* value | Cor. | *p* value |
| Cr2014St01 | $7.21 \times 10^{-7}$ | -0.18 | <0.001 | -0.27 | <0.001 | -0.21 | <0.001 | -0.066 | 0.078 |
| Cr2014St09 | $7.73 \times 10^{-7}$ | 0.14 | <0.001 | -0.27 | <0.001 | -0.057 | 0.15 | -0.019 | 0.63 |
| Cr2057St01 | $6.78 \times 10^{-7}$ | -0.1 | 0.012 | -0.69 | <0.001 | 0.25 | <0.001 | -0.043 | 0.29 |
| Cr2057St09 | $5.79 \times 10^{-7}$ | 0.016 | 0.66 | -0.67 | <0.001 | 0.054 | 0.13 | -0.077 | 0.032 |
| Cr2109St01 | $3.87 \times 10^{-7}$ | -0.39 | <0.001 | -0.56 | <0.001 | 0.53 | <0.001 | -0.19 | <0.001 |
| Cr2109St09 | $3.74 \times 10^{-7}$ | -0.22 | <0.001 | -0.54 | <0.001 | 0.51 | <0.001 | -0.16 | <0.001 |
| Cr2164St01 | $4.14 \times 10^{-7}$ | -0.44 | <0.001 | -0.49 | <0.001 | -0.082 | 0.016 | 0.18 | <0.001 |

Table S3. The proportion of the number of bacteria taxa (ASV) and its relative abundance (% of total reads) present/absent at $T_0$ versus diluted and none-diluted treatments at $T_{12}$ for each experiment (without rarefaction). The diluted predation treatments indicate treatments under 25, 50 and 75% predation dilution factors, whereas the non-diluted predation indicate treatments under the 100% predation dilution factor (full predation effect).

| At $T_0$ | Bacterial taxa **Absent from T0** | | | | Bacterial taxa **Present at T0** | | | |
|---|---|---|---|---|---|---|---|---|
| At $T_{12}$ | Present at **diluted** predation treatments | | Present at **non-diluted** predation treatments | | Absent from **diluted** predation treatments | | Absent from **non-diluted** predation treatments | |
| % of total | ASVs | Reads | ASVs | Reads | ASVs | Reads | ASVs | Reads |
| 2014AprSt1 | 80.91 | 19.01 | 21.24 | 0.89 | 0.12 | <0.001 | 16.11 | 0.40 |
| 2014AprSt9 | 56.90 | 11.15 | 75.86 | 32.11 | 0.57 | <0.001 | 0.48 | 0.02 |
| 2014OctSt1 | 77.97 | 14.49 | 48.92 | 5.40 | 0 | 0 | 8.11 | 0.58 |
| 2014OctSt9 | 78.42 | 9.29 | 62.56 | 11.11 | 0.26 | <0.001 | 0.61 | <0.001 |
| 2015JulSt1 | 61.68 | 5.29 | 41.71 | 2.54 | 0.35 | <0.001 | 5.61 | 0.06 |
| 2015JulSt9 | 80.14 | 18.03 | 39.85 | 4.41 | 0.19 | <0.001 | 4.43 | 0.06 |
| 2016MaySt1 | 75.12 | 7.63 | 39.48 | 2.69 | 0.38 | <0.001 | 5.38 | 0.08 |

Table S4. Environmental and biotic variables in each experiment. Bacterial initial richness was the number of ASV estimated from communities at $T_0$ that rarefied with the lowest number of all samples in this study (at 1598 reads). Bacterial initial density was estimated by the means of bacterial density without dilution (100% predation dilution factor) at $T_0$.

| **Experiment** | **Temperature (°C)** | **$NO_2$+$NO_3$ (µM)** | **$PO_4$ (µM)** | **Initial bacterial richness** | **Initial bacterial density (cells/ml)** |
|---|---|---|---|---|---|
| 2014AprSt1 | 16.86 | 16.41 | 0.46 | 241 | 1592878.56 |
| 2014AprSt9 | 19.57 | 4.15 | 0.28 | 29 | 305721.70 |
| 2014OctSt1 | 26.61 | 13.92 | 0.23 | 171 | 537658.85 |
| 2014OctSt9 | 23.62 | 1.69 | 0.09 | 52 | 314478.76 |
| 2015JulSt1 | 25.48 | 7.29 | 0.58 | 72 | 479109.83 |
| 2015JulSt9 | 26.02 | 0.20 | 0.1 | 220 | 241740.23 |
| 2016MaySt1 | 20.99 | 10.9 | 0.21 | 129 | 724227.43 |

Table S5. Permutation test for analyzing the relationship between growth-based competitiveness versus invulnerability to protist grazing (a) and viral lysis (b), respectively. A null model was generated using the linear regression estimates with randomly shuffled dependent variables (the invulnerability to predation) and repeated 1000 times. The p-value of the Shapiro-Wilk test was estimated to test if the null model was not significantly different from normal distribution (where S-W p-value > 0.05). Z-score and its p-value were estimated to test if the observed value was significantly different from the mean of null model.

| **Experiment** | **S-W p-value** | **z-score** | **p-value** |
|---|---|---|---|
| **(a) Invulnerability to protist grazing** | | | |
| Cr2014St01 | 0.522 | -6.474 | < 0.001 |
| Cr2014St09 | 0.407 | -5.858 | < 0.001 |
| Cr2057St01 | 0.521 | 5.933 | < 0.001 |
| Cr2057St09 | 0.489 | 4.776 | < 0.001 |
| Cr2109St01 | 0.137 | -2.416 | 0.016 |
| Cr2109St09 | 0.039 | -2.090 | 0.037 |
| Cr2164St01 | 0.355 | 1.377 | 0.169 |
| **(b) Invulnerability to viral lysis** | | | |
| Cr2014St01 | 0.628 | -8.794 | < 0.001 |
| Cr2014St09 | 0.666 | -16.798 | < 0.001 |
| Cr2057St01 | 0.337 | -12.034 | < 0.001 |
| Cr2057St09 | 0.955 | -15.250 | < 0.001 |
| Cr2109St01 | 0.704 | -9.154 | < 0.001 |
| Cr2109St09 | 0.591 | -15.348 | < 0.001 |
| Cr2164St01 | 0.844 | -16.144 | < 0.001 |

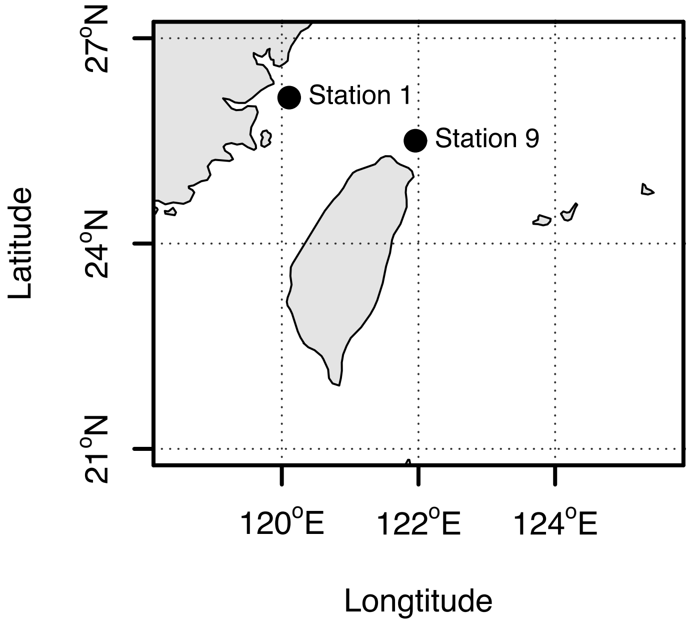


Figure S1. Map illustrating study sites. Samples were taken from the sea surface water and dilution experiments were conducted at Station 1 (April and October 2014, July 2015 and May 2016) and Station 9 (April 2014, July and October 2015).

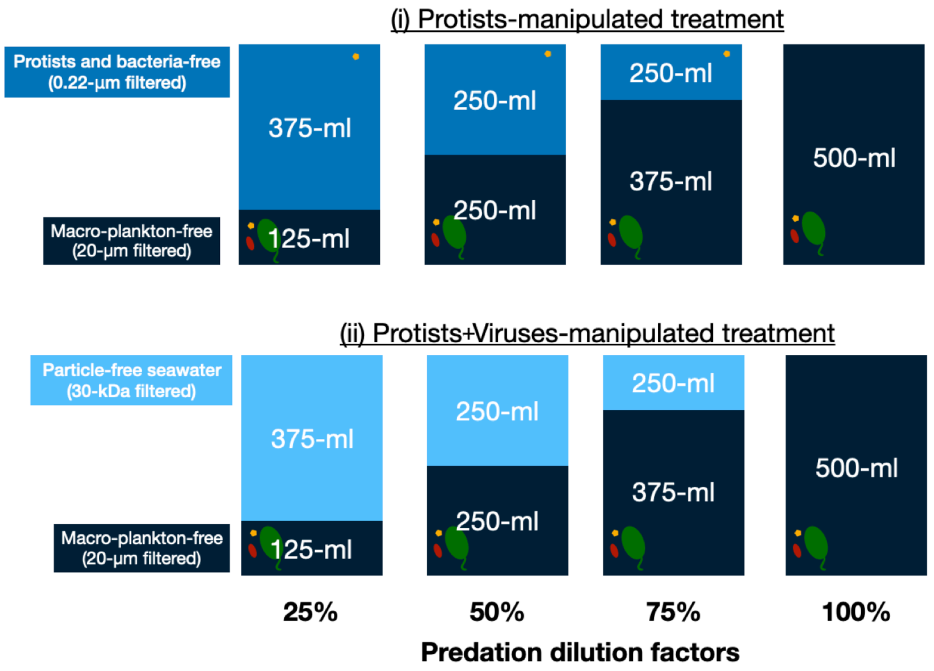


Figure S2. Illustration of predation manipulation experiment. For each set of experiments, three types of seawater were used for creating two combinations of predation manipulation experiments: (i) Protists-manipulated (macro-plankton-free seawater diluted by grazing and bacteria-free water); and (ii) Protists+viruses-manipulated (macro-plankton-free seawater diluted by particle-free seawater). For each combination of predation manipulation, a four-point dilution series was generated, with 25, 50, 75 and 100% of the original predation effect remaining (predation dilution factors). The mixtures were incubated for 12 h in duplicate in sterilized 500-ml polycarbonate bottles at *in situ* temperature in a cabin with water flow taken on board from the sea surface layer and under natural light.

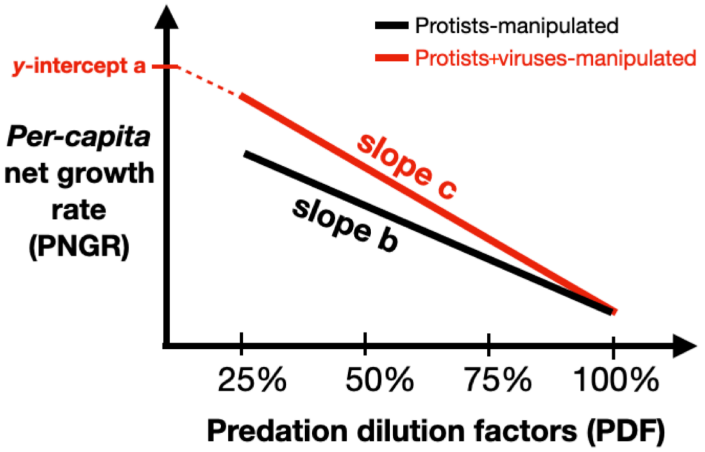


Figure S3. Illustration of relationships between *per-capita* net growth rate (PNGR) versus predation dilution factors (PDF) for each bacterial taxon. Red and black lines represent the PNGR-PDF regression lines under protists- and protists+viruses-manipulated treatment, respectively. ***Bacterial growth-based competitiveness*** was estimated as the *y*-intercept of the PNGR-PDF relationship under protists+viruses-manipulated treatment (*y*-intercept a). ***Bacterial invulnerability to protist grazing*** was the PNGR-PDF regression slope under protists-manipulated treatment (slope b). ***Bacterial invulnerability to viral lysis*** was calculated by subtracting the PNGR-PDF regression slope under protists from regression slope under protists+viruses manipulated treatment (slope c - slope b).

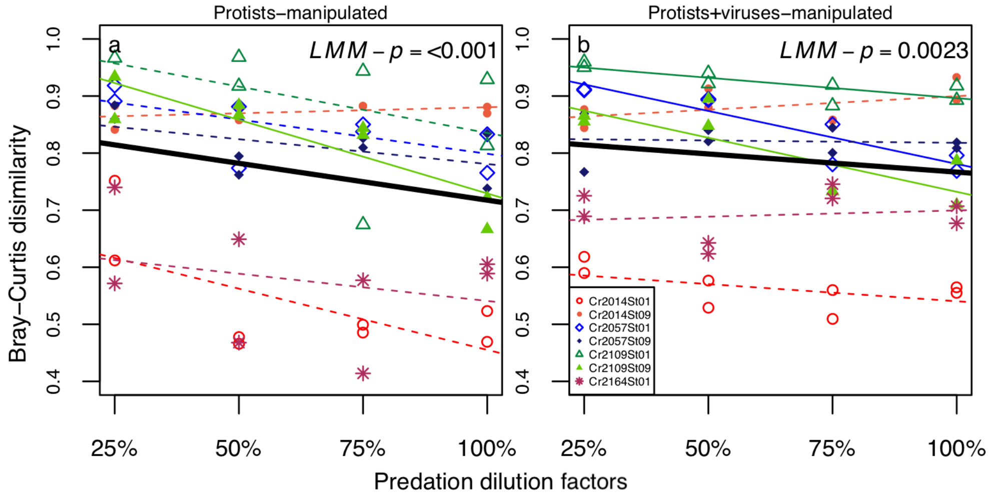


Figure S4. Relationships between predation dilution factors and Bray-Curtis dissimilarity between bacterial community at $T_0$ and $T_{12}$ under (a) Protists-manipulated and (b) Protists-virus manipulated (b) treatment. Colors or symbols indicate different sets of experiments. Solid and dashed lines indicated significant ($p < 0.05$) and non-significant ($p > 0.05$) linear regression. Black bold solid lines represent the significant (*LMM-p* < 0.05) linear regression estimated from linear-mixed effect model with experiments as the random effect.

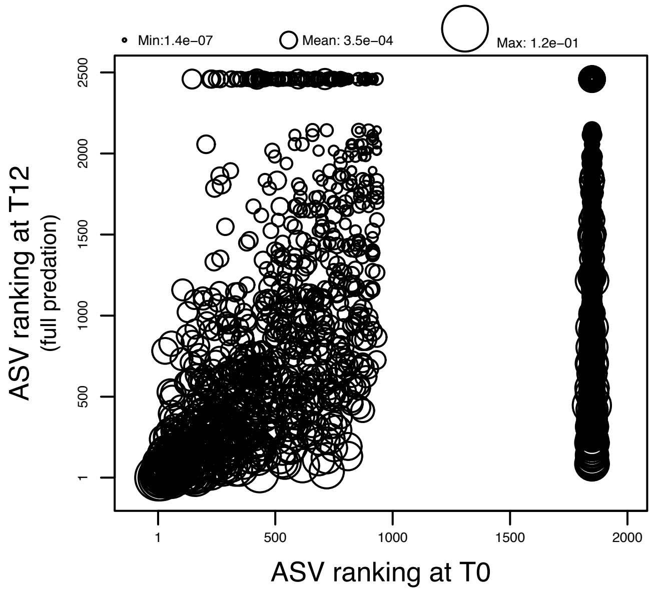


Figure S5. The relationship between bacterial taxa (ASVs) ranking at $T_0$ versus at $T_{12}$ when without predation manipulation (100% predation dilution factor). Point size indicates relative abundance (when non-rarefied and across all experiments) for each ASV.

**Supplementary information**

*Amplification mixture and PCR condition for 16S rRNA gene library preparation*

The V5–V6 region of the 16S rDNA was amplified using the forward primer FIA-787F (5'-[forward index adaptor]-ATTAGATACCCNGGTAG-3') and reverse primer RIA-1046R (5'-[reverse index adaptor]-CGACAGCCATGCANCACCT-3') (Cai *et al.,* 2013). The amplification mix contained: 1 U of *Taq* DNA polymerase (Promega), 1 × reaction buffer, 1.5 mM $MgCl_2$, 0.2 mM dNTPs, 0.2 mM of primers, and 2 ng DNA. The PCR conditions were an initial denaturation at 94 °C for 3 min; 25 cycles of 94 °C for 30 s, 55 °C for 45 s, 72 °C for 1 min; and a final extension at 72 °C for 2 min. Three PCR amplifications were pooled and purified using AMPure XP beads (Beckman Coulter Genomic, CA, USA). Purified products were quantified using a Qubit fluorometer (Invitrogen, Carlsbad, CA, USA) with Qubit dsDNA BR Assay Kit (Life Technologies, USA).

A second PCR was performed using primers containing sample-specific indices and Illumina adaptors. The PCR mix contained: 1 U of *Taq* DNA polymerase (Promega), 1 × reaction buffer, 1.5 mM $MgCl_2$, 0.2 mM dNTPs, 0.2 mM S5 and N7 primers (Nextera Index Kit) and 2 ng DNA purified from the first PCR product. The PCR conditions were an initial denaturation at 94 °C for 3 min; 6 cycles of 94 °C for 30 s, 55 °C for 45 s, 72 °C for 1 min; and a final extension at 72 °C for 2 min. Three PCR amplifications for each sample were pooled and purified using AMPure XP beads according to the manufacturer's instructions. Final products with unique dual-index were quantified using a Qubit fluorometer and pooled in equal concentrations.
